# Multi-strain Analysis of *Pseudomonas putida* Reveals the Metabolic and Genetic Diversity of the Species

**DOI:** 10.1101/2025.11.17.688870

**Authors:** Joshua Mueller, Kalpathy Jayanth Krishnan, Qixing Wei, Ying Hefner, Jonathan M. Monk, Hans Verkler, Juan D. Tibocha-Bonilla, Anthony Ayala, Bernhard O. Palsson, Adam M. Feist, Wei Niu

**Affiliations:** Department of Chemical and Biomolecular Engineering, University of Nebraska-Lincoln, Lincoln, NE, 68588, United States; Department of Bioengineering, University of California, San Diego, La Jolla, CA, 92093, United States; Department of Chemical Engineering, University of Puerto Rico-Mayagüez, Mayagüez, 00680, Puerto Rico; Joint BioEnergy Institute, 5885 Hollis Street, 4th Floor, Emeryville, CA, 94608, United States; Novo Nordisk Foundation Center for Biosustainability, Technical University of Denmark, 2800, Kgs, Lyngby, Denmark; Department of Pediatrics, University of California, San Diego, CA 92093-0760

**Author notes:** Current address: Biological Systems and Engineering Division, Lawrence Berkeley National Laboratory, Berkeley, CA, 94720, USA. These authors contributed equally to this work (Joshua Mueller, Kalpathy Jayanth Krishnan).

**Keywords:** *Pseudomonas putida*, multi-strain analysis, Pan-putida and strain-specific metabolic models, aromatics utilization

## Abstract

*Pseudomonas putida* is a gram-negative bacterial species increasingly utilized in biotechnology due to its robust growth, ability to degrade aromatic compounds, solvent tolerance, and genetic tractability. In this study, we report a comprehensive multi-strain analysis of 164 *P. putida* strains. We performed whole-genome sequencing and hybrid assembly for 40 strains, contributing a ∼8% increase to the available genomic data for *P. putida*. Furthermore, high-throughput phenotypic profiling using the Biolog phenotype microarray system for 24 strains on 190 unique carbon sources, along with 15 aromatic compounds not present on Biolog plates, yielded 4,920 unique strain-phenotype measurements. These data were leveraged to curate GEMs for 24 representative strains, including a refined model for strain KT2440, which comprised 1,480 genes and 2,191 metabolites, achieving a prediction accuracy of 91.2% in carbon utilization. Systematic comparison of genomes and GEMs revealed both conserved core pathways and significant allelic and functional divergence across strains, highlighting strain-specific variation in aromatic degradation. While pathways for protocatechuate and phenylacetate degradation were widely conserved, metabolic capabilities for compounds such as ferulate, phenol, and cresols varied markedly, suggesting adaptation to distinct ecological niches. Alleleome analysis of enzymes such as PcaI and PcaJ revealed distinct, functionally similar clades, indicating possible convergent evolution or horizontal gene transfer. These results provide computable resources and models for selecting *P. putida* strains with desired traits for biomanufacturing and bioremediation and offer insights into the evolution and phylogeny of the *P. putida* species.

## INTRODUCTION

*Pseudomonads* consist of highly adaptable Gram-negative, gamma-proteobacteria with a large range of metabolic capabilities that enables survival in diverse environmental niches^1–3^. Among the over 200 *Pseudomonas* species, *Pseudomonas putid*a (*P. putida*) has been of particular interest for biotechnological applications as a rhizosphere colonizer that generally lacks virulence factors^4–10^. *P. putida* strains have the metabolic versatility to catabolize a wide spectrum of carbon sources, including organic acids, carbohydrates, and aromatic compounds, as well as exhibiting a high tolerance to xenobiotics and resistance to oxidative stress^4–10^. These beneficial traits attracted recent attention for refactoring this species for chemical production^8, 10^ and bioremediation of environmental pollutants^11, 12^. Concurrently, their applicability to industrial biology was propelled by the development of synthetic biology tools to achieve facile and multiplex genetic/genomic manipulations^10, 13–15^. Meanwhile, insights into the metabolic and regulatory networks gained from systems biology studies further accelerate engineering efforts with this species^10, 16, 17^.

Genome-scale metabolic models (GEMs) are becoming indispensable resources in the study and engineering of living organisms^18–22^. However, the GEM of a single microbial strain lacks the content and context needed to investigate the metabolic scope of an entire species. With increasing access to next-generation sequencing, reconstructing the metabolism of multiple strains from the same species becomes possible. Recent efforts on multi-strain genome-scale modeling of bacterial species have shed light on metabolic traits associated with growth adaptation and enabled the delineation of strain subtypes based on metabolic capabilities^23–26^. However, a large-scale multi-strain metabolic analysis of the *P. putida* species is still lacking.

KT2440 is the first completely sequenced^27, 28^, and currently the best-characterized, *P. putida* strain with an FDA HV1 certified status^29^. It is also a major focus of genome-scale modeling and *in silico* simulation studies for the *P. putida* species^30–32^. In this report, we expanded the high-quality *i*JN1463^32^ GEM of the KT2440 strain to develop a pan-genomic metabolic model using genetic content from 164 strains and refined new strain-specific GEMs for 24 strains based on phenotypic data on various substrates. Further analysis revealed large genomic and metabolic diversity within the *P. putida* species that illuminate and should expand current interest in applying *P. putida* for industrial applications.

## RESULTS

### Genome Sequencing and Phylogenetic Analysis

To facilitate multi-strain GEM modeling, we identified 40 under-characterized *P. putida* strains in the American Type Culture Collection (ATCC) for whole genome sequencing (Table S1). A hybrid assembly approach was utilized to generate significantly improved genome assemblies with low contig counts and high N50 values for these strains (Figure 1A & 1B). Only four of the sequenced strains had an N50 less than 1,000,000, with most (72.5%) having an N50 greater than 5,000,000. The average total length of the 40 assembled genomes was 6,056,766 bp. Of the sequenced genomes, 18 were assembled into a single contig, and 15 more had 10 or fewer contigs. This represented a good success rate with 30 of the 40 genomes having high quality assemblies and the remainder still resulting in good quality assemblies except for ATCC 17494.

**Figure 1.**
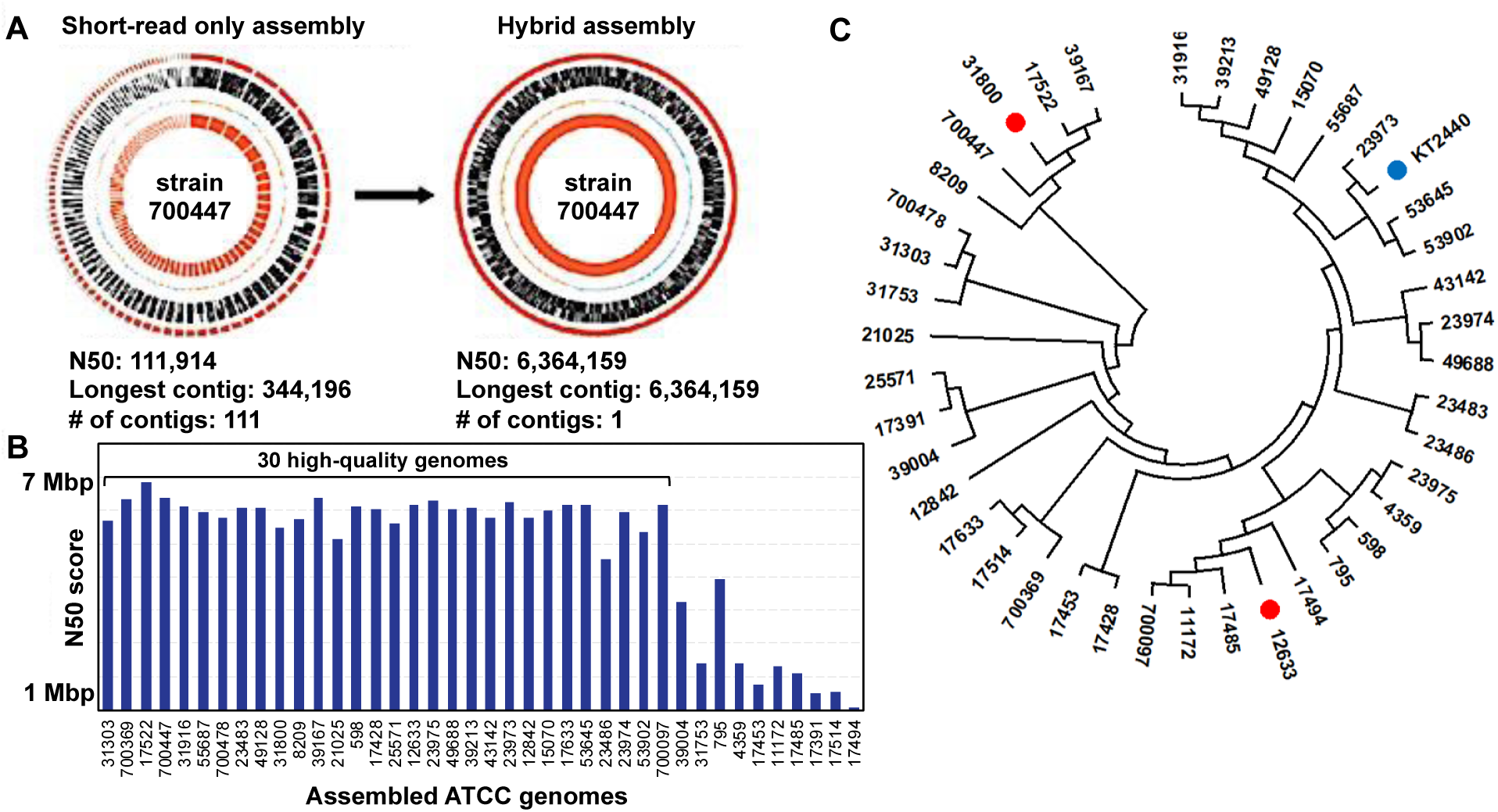
Genome sequencing and phylogenetic analysis of *P. putida* strains. **A.** Comparison of genome completeness between a short-read only assembly (left) and a hybrid assembly approach with the incorporation of long reads (right) for the ATCC 700447 strain, as a representative example. **B.** Comparison of N50 values for assembled genomes of the sequenced strains. Hybrid assembly resulted in significantly higher N50 values that equaled or approached the genome size of assembled high-quality genomes (see Table S1). **C.** Maximum-likelihood tree of the 40 newly sequenced *P. putida* strains. KT2440 is denoted with a blue mark. The two strains chosen for initial reconstruction of the Pan-putida metabolic network are denoted with red marks.

A phylogenetic analysis of the 40 newly sequenced strains together with strain KT2440 was performed. Nine housekeeping genes that had previously been chosen to examine *P. putida* phylogeny were used in a multiple sequence alignment^33^. KT2440 was identified as being phylogenetically distant from many of the newly sequenced strains (Figure 1C). This result indicated that, while *i*JN1463 comprehensively represents the KT2440 strain^32^, this GEM likely does not represent the entire *P. putida* species well. Therefore, a need to build a more expansive metabolic reconstruction to capture the metabolic diversity of the species was identified. Strains ATCC 31800 and ATCC 12633 were chosen in the initial stage of expanding *i*JN1463 into a Pan-putida metabolic network because they are phylogenetically distant from KT2440 (Figure 1C). Due to the relatively smaller number of homologous genes when compared to KT2440, we expected significant increases in new protein sequences and metabolic content by inclusion of the two strains.

### Multi-strain Phenotyping from Growth Analysis on Multiple Substrates

A large body of work detailing the physiology and metabolism of the KT2440 strain enabled the construction of its high-quality GEM^8, 27, 28^. Meanwhile, far less literature exists for other strains of *P. putida*, especially for the 40 under-characterized ATCC strains. This limited primary research data impeded further efforts for generating high quality GEMs of these organisms.

To address the lack of phenotypic data, growth characterization on carbon utilization was conducted for 24 strains that were culturable in chemically defined media, including KT2440, using the Biolog phenotype microarray plates PM01 and PM02A, which contain 190 compounds as sole carbon sources. The Biolog assays enable high-throughput detection of the cellular metabolic activity by continuously monitoring the color change of a cell culture caused by the reduction of a tetrazolium dye^34^. Each strain was phenotypically characterized in duplicates. Of the carbon sources tested, 123 supported detectable metabolic activity in at least one strain of *P. putida* (Table S2). No active respiration was detected with the remaining 67 carbon sources. Furthermore, as 144 of the total 190 compounds had associated <u>Bi</u>ochemically, <u>G</u>enetically and <u>G</u>enomically (BiGG) IDs, which are used as unique identifiers in our GEMs^35^, we were able to use most of the captured Biolog data for metabolic model construction and validation (Figure S1A). Meanwhile, among the 46 Biolog substrates lacking BiGG IDs, 17 supported metabolic activities (Table S2).

Due to the significant interest in aromatic metabolism of *P. putida* for biotechnological applications^36, 37^, we additionally conducted a separate growth characterization of the strains on a selected group of 15 aromatic compounds that were not in Biolog plates (Figure S2). These included lignin-derived aromatics (e.g., *p-*coumarate, ferulate, vanillate, syringate, guaiacol, and syringol) and anthropogenic aromatics (e.g., benzoate, phenol, *m*-cresol, *o*-cresol, *p*-cresol, and terephthalate). *P. putida* strains KT2440, F1, S-16, and B6-2, which had reported growth on several aromatic compounds, were included as positive controls. An agar plate-based assay was used, with each aromatic compound provided as the sole carbon source (Figure S2). The experiment further expanded the structural diversity of carbon substrates and enabled a more in-depth probing of the species’ metabolic capabilities. The obtained data were also incorporated into metabolic model validation and refinement and supplemented the Biolog high-throughput physiology screens, which, in contrast, were based on respiration and not physical colony formation.

To better understand patterns of substrate utilization, the phenotypic data were further analyzed based on the structural features of the compounds. The 190 Biolog substrates were classified into eight groups: (1) organic acids (41 compounds), (2) L-amino acids (21), (3) monosaccharides (17), (4) oligosaccharides (n > 1) (24), (5) sugar acids and alcohols (26), (6) modified sugars (26), (7) aromatics (7), and (8) others (28). The “others” group included D-amino acids, nucleobases, dipeptides, and other miscellaneous compounds. When additional substrates from agar plate experiments were considered, the number of aromatic compounds increased to 22. The metabolic capability of each *P. putida* strain on a given substrate group was quantified as the percentage of compounds within the group that supported metabolic activity. These values were visualized in a clustered heatmap (Figure 2A). To assess group-level trends across the species, a metabolic activity index was defined as the average activity across all strains for each compound group.

**Figure 2.**
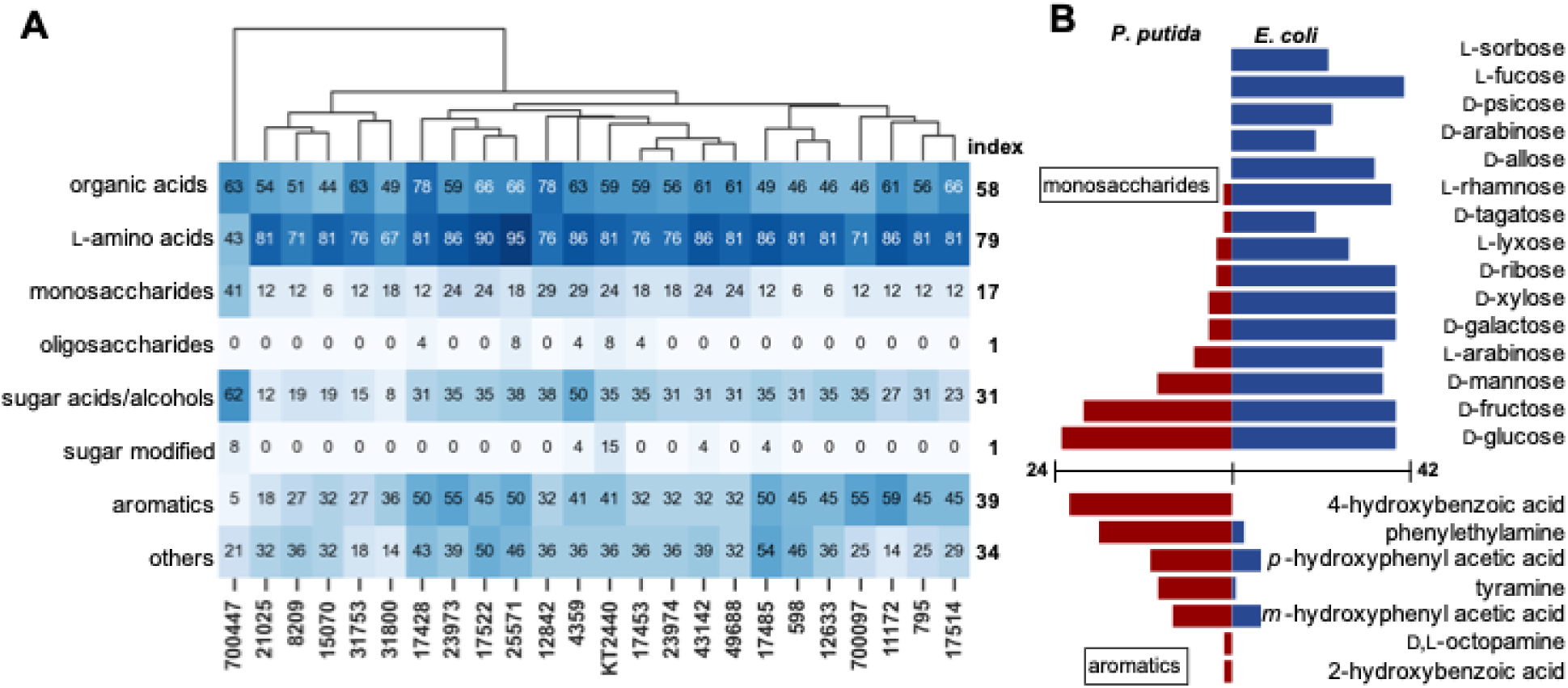
Phenotypic data analysis. (**A**) Clustered heatmap showing metabolic activity profiles of 24 *P. putida* across 205 carbon substrates, including 190 Biolog compounds and 15 additional aromatic compounds. Each row corresponds to a compound group and each column represents a strain. The numerical value within each cell indicates the percentage of compounds in the corresponding group on which the strain exhibited metabolic activity. Color intensity is directly scaled to the numerical values, with darker shades representing a higher percentage. Font color was adjusted to enhance readability. Metabolic activity index values for *P. putida* species on each compound group are shown on the right. (**B**) Comparing metabolic activities of *P. putida* and *E. coli*^38^ on monosaccharides and aromatics. The bar graph illustrates the number of strains that are metabolically active on the indicated substrate. (The graph excludes substrates that do not support the respiration of any strains.).

Among all groups, *P. putida* exhibited the highest metabolic activity on L-amino acids (group 2, index 0.79), with 32% of the compounds supporting respiration in all strains. Notably, every L-amino acid tested supported respiration in at least one strain. Organic acids (group 1, index 0.58) also showed relatively high utilization, with 17% of the compounds supporting metabolic activity in all 24 strains. In contrast, sugar-related compounds generally showed lower metabolic activity index values. None of the strains were metabolically active on 41% of the monosaccharides (group 3, index 0.17), 84% of the oligosaccharides (group 4, index 0.01), and 77% of the modified sugars (group 6, index 0.01). Despite the chemical complexity, the strains demonstrated robust metabolic activity on aromatic compounds (group 7, index 0.39), with only four out of 22 compounds—guaiacol, syringol, syringate, and terephthalate—failing to support respiration. Interestingly, clustering analysis revealed that strain ATCC 700447 was metabolically distinct from the other 23 strains. This strain exhibited higher activity on monosaccharides and sugar acids/alcohols, but lower activity on L-amino acids and aromatic compounds, suggesting unique metabolic preferences or regulatory traits.

We conducted a parallel analysis using previously reported Biolog phenotyping data from 42 *E. coli* strains (Figure S3)^38^. In contrast to *P. putida*, *E. coli* exhibited strong metabolic activity on sugars and their derivatives, but showed limited activity on L-amino acids, organic acids, and aromatic compounds. These differences in substrate utilization are particularly evident for monosaccharides and aromatics, as illustrated in Figure 2B. Monosaccharides, which are commonly used as starting materials in physiological studies and biomanufacturing, were poorly utilized by *P. putida*. Among the 17 simple sugars tested, only D-glucose (23 strains) and D-fructose (20 strains) supported respiration in the majority of *P. putida* strains. Other common sugars such as D-mannose, L-arabinose, D-xylose, and D-galactose supported growth in only 10, 5, 3, and 3 strains, respectively. Notably, 7 out of 17 monosaccharides did not support respiration in any *P. putida* strain. In contrast, only two monosaccharides (i.e., L-glucose and D-fucose) failed to support metabolic activity in *E. coli*, which demonstrated broad utilization of both C5 and C6 sugars. This trend was reversed when aromatic compounds were provided as the sole carbon source in Biolog assays. All tested aromatic substrates supported respiration in at least one *P. putida* strain, whereas four out of seven did not support any metabolic activity in *E. coli*. These findings underscore fundamental phenotypic differences in substrate preferences between the two species, reflecting their distinct ecological niches and evolutionary adaptations.

### Analysis of Genomic Content

The expanded genome set of *P. putida* allowed for the generation of an elementary pan-genome and further analysis into the core metabolic genome of *P. putida.* The Cluster Database at High Identity with Tolerance, CD-HIT^39^ program was used to identify orthologous gene families/clusters across 164 strains, which resulted in a ‘pan-genome’ with a core gene family set of 1,641 genes (Materials and Methods). The size of the core genome was plotted by ordering strains based on the number of genes shared with the pan genome (Figure 3A). This approach differs from previous analyses where the strain order was randomized, and an average of the core-genome size was plotted after numerous randomizations^25, 40^. As a result, this core genome size plot is more suitable for analyzing the phylogenetic relationships of the strains (Figure 3A). More distantly related strains result in a steeper line as the core-genome decreases in size more rapidly. For a group of closely related strains, once a set of core-defining characteristics is reached, the core genome size is expected to approach a defined value, even as more strains are added. However, the size of the core genome of the selected *P. putida* strains showed an accelerated reduction after initially appearing to stabilize when approximately 100 strains were included.

**Figure 3.**
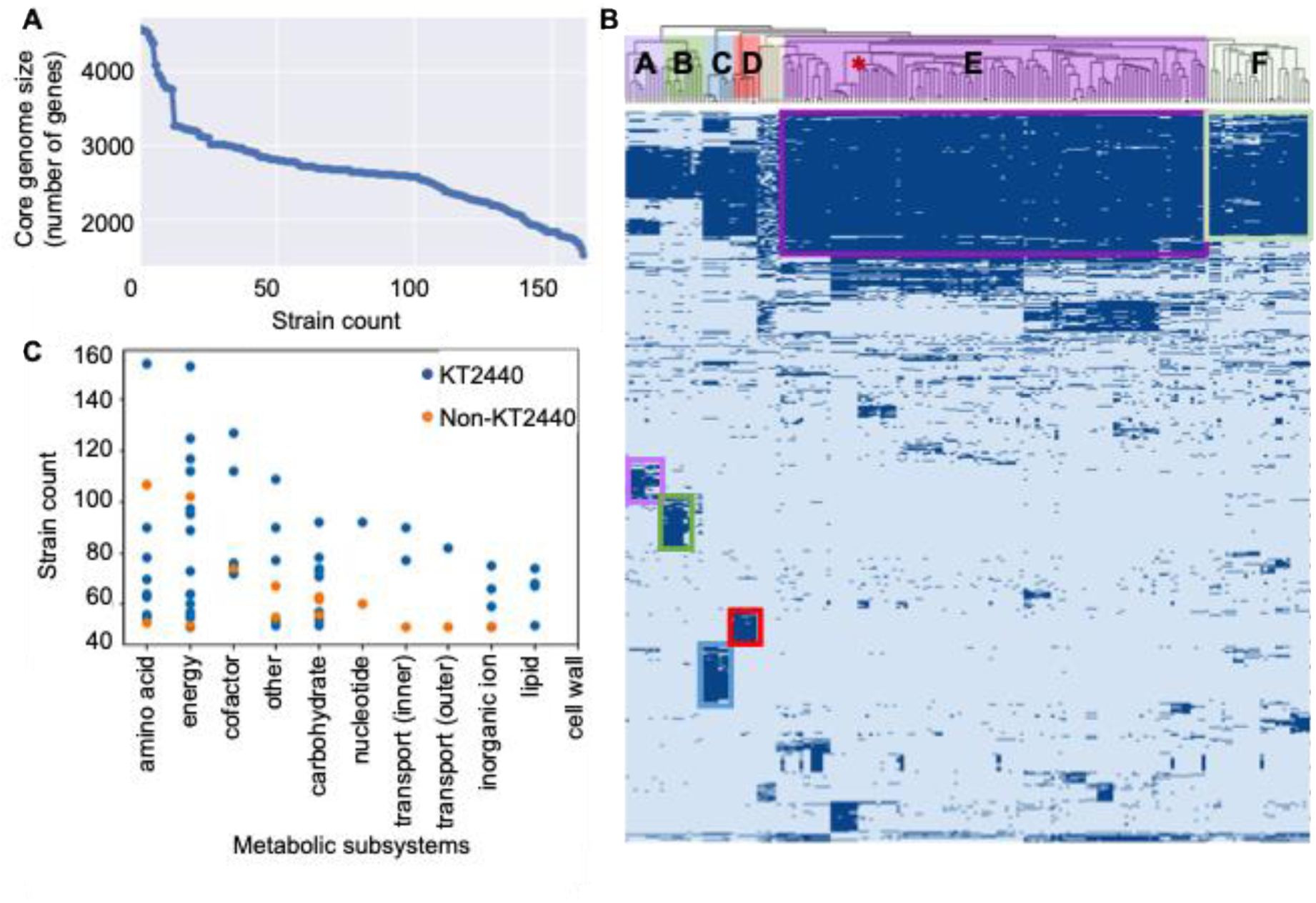
Genomic and allelic analysis. **A.** Core genome size plot. The first data point represents the initial “core” genome defined by strain ATCC 23973. The core genome was refined with the inclusion of additional strains. **B.** CD-HIT gene family clustering based on amino acid sequences. Strains are grouped based on the degree of genetic similarity. Each column represents a strain, and each row represents a gene family predicted by CD-HIT. The color indicates the presence (dark blue) or absence (light blue) of a gene family. Strains were grouped into Blocks A to F, with the position of strain KT2440 marked by a red asterisk. **C.** Frequency of specific genomic alleles across all 164 strains of *P. putida*. Each dot represents a specific gene, grouped by metabolic subsystems. Blue dots correspond to alleles in KT2440. Orange dots are alleles absent in KT2440 but are found in multiple other *P. putida* strains. Strain counts of alleles are represented on the y-axis.

We further probed the genomic diversity by assigning ^39^the genes from all the 164 strains into 108,745 gene families based on the amino acid sequences. Strains were then clustered based on shared and unique gene families to identify genomic diversity. A total of six strain clusters were revealed (Figure 3B) using a hierarchical clustering method. Blocks E and F harbored the majority of the studied strains, which shared significant genetic content. Strains in block E were more tightly related, and GEMs of strains in this block had slightly higher accuracy for growth predictions (92%) in comparison to the average accuracy for all strains (91%). Further investigation focused on identifying any patterns or potential reasons associated with dissimilarities of strains clustered in Blocks A to D. Each cluster had a set of unique genetic features consisting of hundreds of gene families (marked with colored boxes in Figure 3B). Strains in blocks C and D were likely clustered due to the identical or similar source of isolation from a single study (Table S6). These are likely closely related strains that had little time for evolutionary divergence. Meanwhile, strains in blocks A and B were isolated from diverse environmental sources in several unrelated studies (Table S6), indicating true genomic similarity following separate evolutionary events. These strains had the lowest number of common gene families with strains in block E and potentially represent different *Pseudomonads* species or subspecies from *P. putida.*^41^

We next compiled protein sequence-based alleleomes for each gene family to investigate genetic conservation and variation across the *P. putida* species^42^. Each allele was defined as a unique amino acid sequence, while allele frequency was assessed by counting the number of strains harboring the same allele. Table S8 summarized the top alleles, defined as those present in over 50% of the strains analyzed. These high-frequency alleles are predominantly associated with essential cellular functions, such as ribosomal subunits in translation and RNA polymerase subunits in transcription. Notably, protein size was found to be a contributing factor to allele frequency. High-frequency alleles tended to be smaller, with an average length of 138 amino acids (median: 118 a.a.), compared to the overall average protein size in the pangenome (332 a.a., median: 284 a.a.). Nonetheless, several large proteins, including the 30S ribosomal protein S1 and transcription terminator Rho, also exhibited high allele frequency, indicating strong evolutionary conservation.

Among metabolic proteins, IlvH (163 a.a.), the small subunit of acetolactate synthase, emerged as the most conserved, reflecting the importance of branched-chain amino acid biosynthesis. We also identified PaaB (93 a.a.), involved in phenylacetate degradation, and two key translational regulators of carbon metabolism-CsrA (62 a.a.) and Crc (259 a.a.)^43–46^-as highly conserved across strains (Table S8). Despite the presence of multiple alleles, especially for the larger Crc protein (31 total), these variants mainly consist of conservative substitutions (Figure S5), where amino acids with similar biochemical properties are exchanged, likely preserving protein structure and function. The strong sequence conservation suggests selective pressure to maintain core regulatory function. Together, these findings point to a robust set of conserved metabolic and regulatory elements across the species. This alleleome analysis not only highlights genes likely under strong purifying selection but also serves as a basis for pinpointing variable loci suitable for strain-specific engineering.

Further genomic comparison across all 164 *P. putida* strains, in conjunction with the generated metabolic models, enabled a systematic evaluation of metabolic subsystems based on gene conservation and reaction content across strains. The analysis identified Amino Acid Metabolism, Energy Production, and Cofactor and Prosthetic Group Metabolism as the most conserved subsystems, underscoring their essential roles in cellular maintenance and growth. (Figure 3C). In contrast, Cell Membrane Metabolism was the only subsystem lacking identical alleles in at least 50 strains, consistent with previous hypothesis that links membrane protein variability to environmental adaptation^47, 48^. These results provide a valuable foundation for strain selection and engineering, as they highlight both highly conserved metabolic functions and strain-specific adaptive traits that can influence industrial performance, such as tolerance, transport, and biofilm formation.

### Updates to metabolic network reconstruction and strain specific models

The Pan-putida metabolic reconstruction was built based on the genome sequences of 164 strains, including KT2440, 40 sequenced ATCC strains from this study, and 123 strains with either whole genome or completed genome status from the Pathosystems Resource Integration Center (PATRIC) database (Table S3)^39, 40^. Phenotypic data from Biolog plates and agar plate assays, as described earlier, were used to validate and ‘gap-fill’ 23 strain specific models which were genereated using bi-directional blast to identify orthologous protein families. (Materials and Methods). These gapfilled reactions were added to the metabolic reconstruction resulting in a expanded pangenome model including a total of 2,326 genes and 3,301 reactions, encompassing 2,525 metabolites. This represents increases of 59% (864), 13% (374), and 17% (372), respectively, compared to *i*JN1463^32^ (Table 1) enabling a more comprehensive analysis of the metabolic diversity of *P. putida* species.

**Table 1.** Properties of the pangenome-wide model compared to iJN1463.

|  |  | **<u>This study</u>** | **<u>iJN1463</u>** |
| --- | --- | --- | --- |
| **Genes** | Total | 2326 | 1462 |
|  | Metabolic | 1810 | 1014 |
|  | Transport | 531 | 461 |
| **Metabolites** | Total unique | 2525 | 2153 |
|  | Cytoplasmic | 1527 | 1339 |
|  | Periplasmic | 496 | 465 |
|  | Extracellular | 407 | 349 |
| **Reactions** | Total | 3301 | 2927 |
|  | Cytoplasmic | 1801 | 1604 |
|  | Periplasmic | 133 | 129 |
|  | Extracellular | 423 | 368 |
| **Transport reactions** | Total | 915 | 826 |
|  | Cytoplasm to Periplasm | 486 | 456 |
|  | Periplasm to extracellular | 337 | 309 |
|  | Cytoplasm to extracellular | 40 | 10 |

Subsystem classification of reactions highlights significant additions to transport reactions (16% addition), central carbon (61%), amino acid (28%), alternative carbon (56%) and aromatic carbon (16%) metabolism showcasing significant additions of new catabolic capabilities, previously not represented in P. putida species (Figure 4 A).

**Figure 4.**
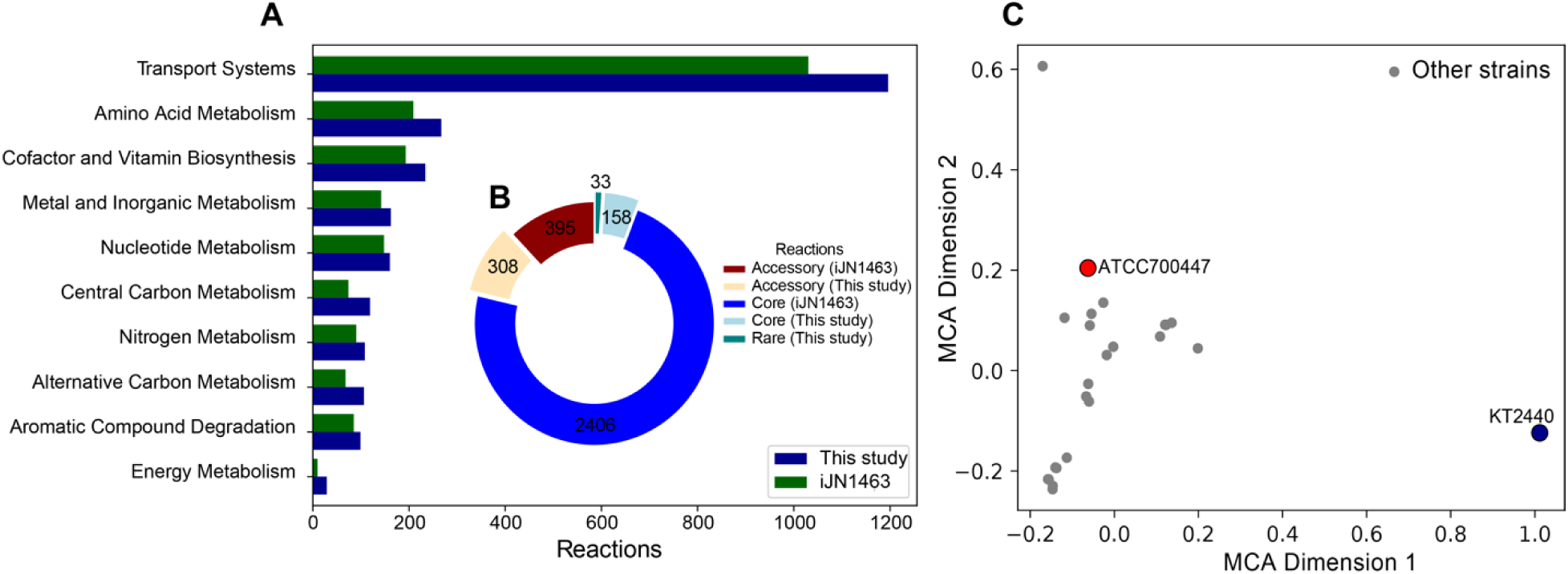
Updates to the metabolic network construction of P. putida organized by. (**A**) Reaction subsystem (**B**) Pangenomic classification (Core reactions are defined as reactions in 98% of strains, accessory reactions are found in 10-98% of strains and rare reactions are found in <10% of strains. (**C**) Multiple Correspondance Analysis scatter plot of 24 strain specific metabolic models with model strain KT2440 and strain ATCC700447 highlighted.

The presence/absence of reactions in the updated reconstruction and iJN1463 across these 24 strain specific GEMs were used to calculate ‘core’ and ‘rare’ reactions, found in >98% of the strains or <10% of strains. Intermediately present reactions were classified as ‘accessory’ reactions. The updated reconstruction provides 158 additional core reactions, 308 accessory reactions and 33 rare reactions (Figure 4 B), thereby providing a better template to construct strain specific *P. putida* models. Multiple correspondence analysis of the reaction presence/absence matrix across the strains revealed that several strain specific GEMs had significantly different network structure compared to the model KT2440 network. Strain ATCC 700447 highlighted for example, is phylogenetically (Figure 1C) and metabolically distant (Figure 4C) from KT2440 and has a strong capacity for utilizing sugars and sugar alcohols but is less capable of using aromatics as demonstrated in the phenotyping experiment (Figure 2A), indicating agreement between the metabolic reconstruction and experimental data. This highlights a significant update to our knowledge of metabolic network and related capabilities in *P. putida* species.

### Analysis of Aromatic Utilization Capabilities

Microbial degradation of aromatics relies on numerous peripheral reactions and pathways, as well as a few key central pathways, to feed structurally diverse carbon sources into central metabolism. ^49^Using bi-directional blast on the gene protein reaction rules from the updated reconstruction ^48^ (Materials and Methods), we generated metabolic models predict the growth capabilities of all 164 *P. putida* strains on 17 aromatic compounds (Figure 5A). With few exceptions, nearly all the strain models were able to metabolize the aromatic amino acids, L-tyrosine (162) and L-phenylalanine (161). Other aromatics that supported the growth of most models included protocatechuate (150), benzoate (129), and phenylacetate (129). The first two can be metabolized through either the protocatechuate or the catechol branch (*ortho*-cleavage) of the *b*-ketoadipate pathway (Figure 5B), and phenylacetate can be metabolized through the aerobic phenylacetyl-CoA pathway (Figure 5B)^11^. Because the utilization of 4-hydroxybenzoate, vanillate, and phenylethylamine requires simple extentions of the above two major pathways, i.e., a hydroxylase or a demethylase to convert 4-hydroxybenzoate or vanillate to protocatechuate and a deaminase plus an aldehyde dehydrogenase to convert phenylethylamine to phenylacetate (Figure 5B), these three substrates also supported the *in silico* growth of the majority of the strains (Figure 5A). We also observed a stronger ability to utilize coumarate (59) and ferulate (57), which can be derived from lignin degradation, compared to various phenolic compounds, including phenol (22) and *o-*/*m-*/*p*-cresols (10/9/9), which are considered environmental pollutants (Figure 5A). Furthermore, the metabolic capabilities for these two groups of compounds rarely co-exist in the same strain, suggesting possible distinct growth niches.

**Figure 5.**
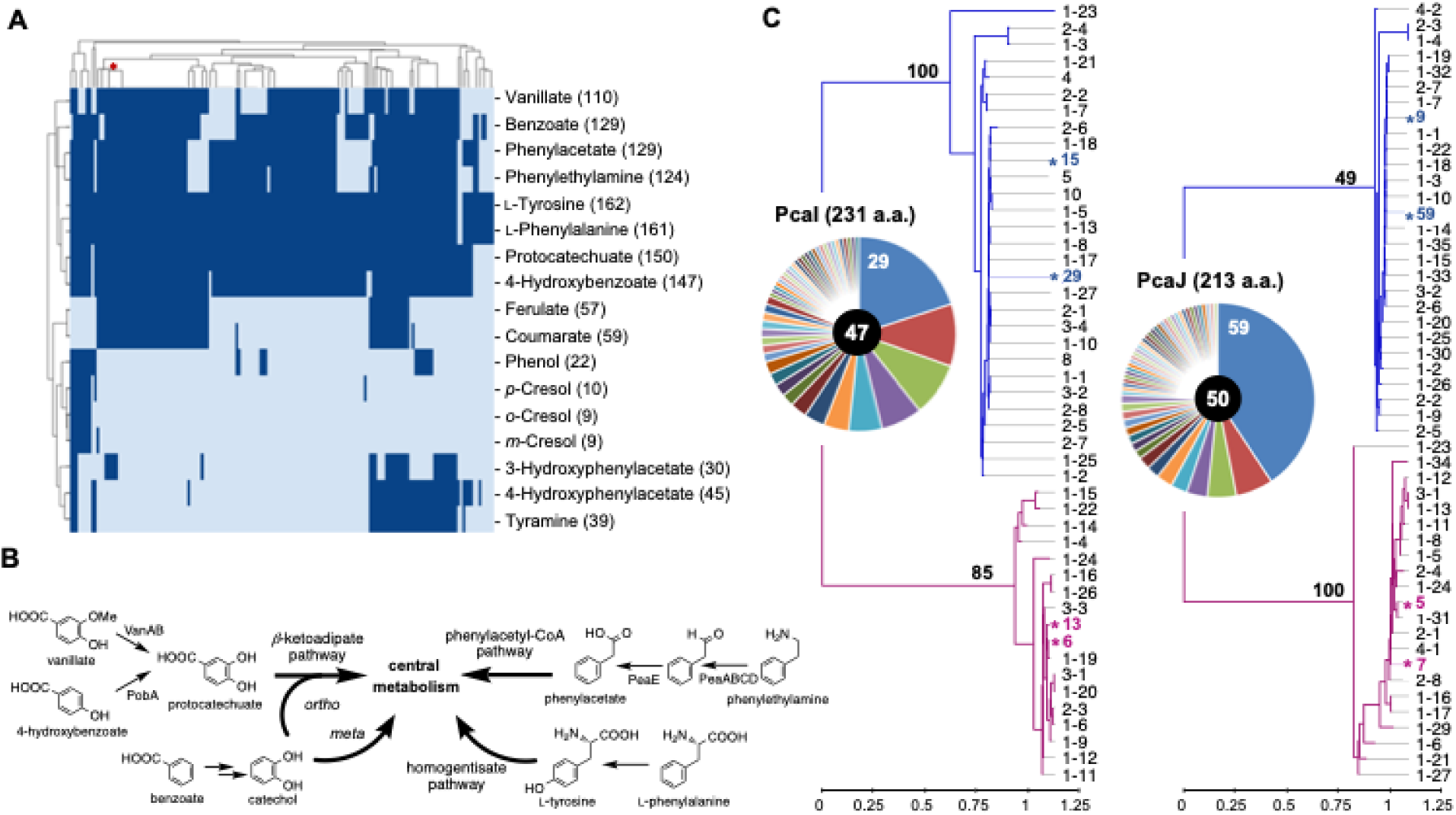
Aromatics utilization by *P. putida* strains. **A.** Clustering of *P. putida* strains by model-predicted growth on aromatic compounds. The position of strain KT2440 is marked with a red asterisk. Values in parenthesis indicate the predicted number of strains with growth on the specified carbon source. **B.** Metabolism of preferred aromatic compounds by *P. putida*. **C.**Analysis of the PcaI and PcaJ alleleomes. Alleleomes are plotted as pie charts based on the strain count of each allele, with values for the dominant alleles marked. Allele counts are shown at the center of the charts. The phylogeny of PcaI and PcaJ was analyzed using the maximum likelihood method. Major clades are shown in blue, and minor clades are shown in purple. Bootstrap values for the main nodes in the two clades are shown in bold. Alleles are named according to their strain counts. The corresponding strains are listed in Table S9. Dominant alleles are marked with asterisks. Scale bars represent the number of substitutions per site.

We conducted a comprehensive alleleome analysis of enzymes involved in two central pathways, the *b*-ketoadipate and the phenylacetyl-CoA pathways, as well as the peripheral benzoate degradation pathway, due to its prevelance in *P. putida* species (Figure S6). The composition of 27 alleleomes was visualized as pie charts with segments representing the strain count of each allele (Figure S7). The majority of the alleleomes had a single dominant allele, while a few consisted of two top alleles with the same strain count. Variations in the allele count indicated varied degrees of conservation between the pathways. The protocatechuate branch of the *b*-ketoadipate pathway was more conserved than the catechol branch. Corresponding allelomes of pathway enzymes had an overall lower allele count, and the dominant alleles had higher strain counts. Meanwhile, the phenylacetyl-CoA pathway showed a surprisingly low overall degree of conservation, even though it contains the most conserved protein, PaaB (Figure S7B), which had a dominant allele in 94 strains and an allele count of 14. Most enzymes in the rest of the pathway had alleles identified in fewer than ten strains, with allele counts close to or exceeding 70 (Figure S7B). The peripheral pathway for benzoate to catechol conversion showed an intermediate degree of conservation, with allele counts lower than 60 for most of the steps.

To further understand the evolution of each alleleome, the alleles were subjected to phylogenetic analysis using the maximum likelihood method, treating each allele as a single sequence regarless of its strain count (Figure S8). The majority of the alleleomes formed a major clade and a minor clade. Branch lengths of clades were generally smaller than 0.1 substitutions per site. Similar values were observed for the node braches. On average, 85% of the alleles, including the dominant allele(s), fell into the major clade. The results indicated that most of the alleleomes were formed by alleles of high sequence homology, though a few exceptions were noted. Clades of the *a* (PcaG) and the *b* (PcaH) subunits of protocatechuate 3,4-dioxygenase showed significantly longer branch lengths ranging from 0.85 to 3.2 substitutions per site (Figure S8A). Small clades of each alleleome consisted of three alleles belonging to the same three *P. putida* strains (Table S9). A more noticable deviation was observed for alleleomes of the two subunits of 3-oxoadipate CoA-transferase (PcaI and PcaJ) (Figure 4C). In addition to having longer-than-average branch lengths, only around 60% of the alleles formed the major clades. Alleles with high strain counts were also observed in the minor clade, representing alleles from a total of 40 strains (Table S9).

## DISCUSSION

*Pseudomonas putida* species can degrade diverse aromatic compounds, tolerate solvents and oxidative stress, making them valuable hosts for industrial bioproduction, bioremediation, and lignin valorization. However, much of the current metabolic knowledge derives from the KT2440 strain. By integrating genome-scale metabolic reconstructions (GEMs) with pangenomic and phenotypic data, we expanded metabolic insights across the species and revealed notable variation in aromatic carbon utilization (e.g., phenol, cresols). Strain-specific GEMs generated from this analysis provide predictive tools for metabolic engineering.

Since the start of this study, the number of publicly available *P. putida* genomes has grown from 123 to over 540 and 444 assemblies on NCBI and BV-BRC^50^, respectively, reflecting rising interest in the species. The 40 newly sequenced genomes reported here increase the species’ diversity by ∼8% and broaden its ecological representation. Our pan-*P. putida* reconstruction, based on 164 globally distributed strains, captures metabolic diversity beyond KT2440 and offers a foundation for future comparative studies. Using the pan-genome, we built and validated 24 strain-specific GEMs with Biolog and growth assays. These models showed high predictive accuracy, while false negatives point to latent or uncharacterized pathways worth exploring through adaptive evolution^51, 52^. Forty-six substrates, including polymers (e.g., laminarin) and anthropogenic compounds (e.g., bromosuccinate), lacked sufficient annotation, underscoring the need for improved biochemical data to enhance model completeness. The refined KT2440 GEM (1,480 genes, 2,995 reactions) improved growth prediction accuracy by 3.8% across 159 substrates, reinforcing its utility for systems biology. Additionally, comparative analyses with *E. coli* further highlighted *P. putida*’s superior catabolism of aromatics and organic acids, consistent with its ecological niche.

Phenotyping results showed 74% agreement between Biolog and plate assays. Discrepancies stem partly from Biolog’s redox-based detection, which can register metabolic activity without biomass formation as illustrated by formate oxidation without assimilation in KT2440^53, 54^. This phenomenon was also observed in 17 additional strains. Given reports of multiple formate dehydrogenases in KT2440^55^, the finding suggests further investigation into substrates that support energy metabolism without contributing to biomass production.

The metabolic profile of *P. putida* reflects its rhizobacterial ecology, as the β-ketoadipate pathway, essential for aerobic aromatic degradation of plant polymers and soil pollutants, is widespread across strains^56, 57^. Both protocatechuate (150 strains) and catechol (129 strains) branches were present in most of the 164 analyzed genomes. Approximately 78% of the strains also carried the phenylacetyl-CoA pathway, which metabolizes distinct aromatic substrates such as amino and phenylalkanoic acids^58^. While many microbes convert L-phenylalanine via phenylacetate, *P. putida* instead employs the homogentisate route^59^. In contrast to the highly conserved ortho cleavage metabolism for protocatechuate, other peripheral aromatic pathways were diverse. The meta-cleavage route of catechol occurred in ∼10% of strains, and ortho-/meta-cleavage of dihydroxyphenylacetate and dihydroxytoluenes involved in tyramine/hydroxyphenylacetate^60, 61^ and o-/m-/p-cresol^62–64^ degradation were also detected. Alleleome and phylogenetic analyses revealed conserved enzyme families with few localized divergence. For example, PcaI and PcaJ enzymes each forming two clades across 60% and 40% of strains, suggesting evolutionary divergence or horizontal gene transfer. Despite variation in strain counts and protein size, these alleleomes provide a framework for linking sequence diversity to enzyme function in future strain engineering.

Both experimental and in silico analyses confirmed limited sugar metabolism across *P. putida* species, consistent with prior efforts to expand KT2440’s sugar utilization^65–70^. A few strains capable of C5/C6 sugar catabolism were identified in this study and offer potential alternative engineering chassis. In addition, the inefficient utilization of syringyl lignin derivatives, observed both experimentally and genomically^71–73^, together with the lack of aromatic oligomer degradation function, pinpoints metabolic bottlenecks that must be addressed to enhance the species’ capacity for lignin valorization.

## MATERIALS AND METHODS

### Media Preparation

All commercial chemicals are of reagent grade or higher. All solutions were prepared in deionized water that was further treated by Barnstead Nanopure® ultrapure water purification system (Thermo Fisher Scientific Inc). Preparation of LB media and M9 salts followed reported recipes^74^. M9 media contained MgSO_4_ (0.12 g/L), CaCl_2_ (0.028 g/L), and trace metal solution^75^, and carbon sources at indicated concentrations. The pH values of all minimal media were adjusted to 7.0. Media containing syringate and syringol were also supplemented with cysteine (6 mM) to prevent oxidation.

### Strain Acquisition and Sequencing

Forty strains that are classified as *Pseudomonas putida* and have no publicly available data of the whole genome sequence were purchased from the ATCC (Table S1). Additional three *P. putida* strains, i.e., S-16, B6-2, and F1, were purchased as controls in growth experiments. Received strains were recovered following the instructions by ATCC. Genomic DNA (gDNA) of the 40 strains was extracted from overnight cultures grown in 3 mL of LB using the QIAamp DNA Mini Kit (QIAGEN). For each strain, gDNA libraries were prepared and sequenced using both the MinION (Oxford Nanopore Technologies) and Illumina HiSeq platforms. Briefly, a library for MinION device was prepared using the ONT Rapid Barcoding Sequencing kit (SQK-RBK004) according to the manufacturer’s protocol with the following modifications: to two separate 0.2 mL PCR tubes, 1 and 0.5 μg gDNA were diluted to 9 μL in ONT EB buffer (10 mM Tris, 50 mM NaCl, pH 8.0). The barcoded fragmentation mix was added in a ratio of 3:1 and 1:1 (μg gDNA: μL fragmentation mix) to the 1 and 0.5 μg samples, respectively. The total library volume was brought to 75 μL with the ONT EB buffer and half the library was loaded onto the MinION R9.4 flow cell without loading beads. Sequencing was performed for 6 h. The FAST5 files were basecalled using the ONT Guppy basecaller (v3.2.2). Quality filtering was enabled with default settings using high accuracy, high accuracy with base pair modification, and fast base calling algorithms. Basecalled reads were demultiplexed with qcat (v1.1.0) prior to assembly. Libraries for the HiSeq system were generated using the TruSeq DNA sample preparation kit (Illumina Inc.) following standard protocols. The libraries were sequenced with a paired-end protocol and read lengths of 150 nucleotides. Whole-genome sequences were hybrid assembled using a combination of the Illumina short reads and the MinION long reads using Unicycler (v0.4.9)^76^. Genomes were annotated using Prokka (v1.12)^77^. BUSCO^78^ and QUAST^79^ were utilized to assess quality and completeness of the genome assemblies. The generated report is deposited in https://doi.org/10.5281/zenodo.17382094 and all assembly fasta files have been submitted to NCBI under the BioProject ID PRJNA1347105.

### Phylogenetic Analysis of *P. putida* Strains

Nine housekeeping genes, including *argS, dnaN, dnaQ, era, gltA, gyrB, ppnK, rpoB, and rpoD*^33^, were used to generate a maximum likelihood tree. The nucleotide sequences of these genes from KT2440 and the 40 newly sequenced strains of *P. putida* were concatenated and aligned using the MUSCLE^80^ algorithm through the MEGA-X software^81^. The evolutionary history was inferred using the Maximum Likelihood method and Tamura-Nei model^82^. The tree with the highest log likelihood (–98751.79) was selected. Initial tree(s) for the heuristic search were obtained automatically by applying Neighbor-Join and BioNJ algorithms to a matrix of pairwise distances estimated using the Maximum Composite Likelihood (MCL) approach, and then selecting the topology with superior log likelihood value. Codon positions included were 1st+2nd+3rd+Noncoding. There was a total of 15175 positions in the final dataset. The phylogenetic tree construction was performed using the MEGA-X software^81^.

### Phenotypic Characterizations

For Biolog growth tests, the targeted strains were grown overnight at 30 °C with shaking in M9 minimal medium containing glucose (4 g/L). For strains with poor initial growth, including ATCC 598, 795, 17421, 17428, 17485, and 17494, multiple passages in the minimal media were performed. LB media was supplemented to a few strains, including ATCC 13795, 17502, and 700447. The overnight cultures were used to seed the Biolog plates PM1 and PM2A, which were then placed in the OmniLog® microplate reader and ran for 48 hours at 30 °C without shaking. All strains were cultured in triplicate plates. To convert opacity respiration readings into growth curves, the signal from each well was processed through a Savitzky-Golay filter to smooth the data. The filter used a window length of 50 and a polynomial degree of 3. The maximum signal value was recorded, and a control group was formed from the negative control wells. A one-sided z-test was performed to calculate the p-values for each well, which were corrected for multiple hypothesis tests using the Bonferroni correction. If the adjusted p-value was below 0.05, the well was considered to exhibit a significant growth signal and was classified as showing growth; otherwise, no growth was assumed.

For non-Biolog growth tests, the seed culture of *P. putida* strains was started in LB media at 1 mL scale in 96-well deep well plates. Following overnight cultivation at 30 °C and 250 rpm, a small volume of the seed was transferred to the surface of a M9 agar plates containing the indicated carbon source (0.25%, w/v) using a stainless steel 96-head pin tool. Duplicate plates were prepared for each carbon source. The plates were incubated at 30 °C for 72 h. Cell growth was documented using a Bio-Rad ChemiDoc imaging system. All Biolog phenotype microarray data is available for visualization and parsing on PMKbase.com^38^. Additionally, all phenotypic microarray data is also available on https://doi.org/10.5281/zenodo.17382094.

### Pan-putida Model Construction

The Pan-putida metabolic reconstruction was built based on the genome sequences of 164 strains, including KT2440, 40 sequenced ATCC strains from this study, and 123 strains with either whole genome or completed genome status from the BV BRC database (Table S3)^83, 84^. *i*JN1463, the latest curated metabolic reconstruction for P. putida KT2440 along with obtained genomic and growth data was used to create a draft Pan-putida metabolic model by following a reported procedure^85^. Briefly, the genomes of strains ATCC 12633 and 31800 were used to enrich metabolic contents at the initial stage of the model development. Based on the annotated genomes, we first manually identified genes which had associated Enzyme Commission (EC) numbers that corresponded to a specific metabolic reaction. Identified reactions were then added to the Pan-putida model and the supporting genetic information was added to a Pan-*putida pan-*genome. The process resulted in the first draft of the pan-genome model. At the second stage, the draft was revised during the validation process by incorporating metabolic data from growth analysis (Figure S1). Comparison between the real growth data and *in silico* prediction revealed gaps between strains’ metabolism and model contents. This information was used to search for genomic content that could reconcile the discrepancies. We attempted gap-filling by searching the BiGG database for matching metabolic step(s) in previously modeled pathways. The underlying genomic content was used to identify homologs in *P. putida* strains. In the case when no pathway had been previously modeled, the reactions were manually added along with supporting genomic content. The updated metabolic model and strain specific models can be found on https://doi.org/10.5281/zenodo.17382094, along with subsystem annotations for each reaction in csv format since these annotations can get lost in reading/writing models.

### Strain-specific Model Construction

The Pan-putida model was used to create strain-specific GEMs^85^. The best Bi-directional BLAST Hit (BBH)^86^ analysis was done to find homologues between individual strains and the pan-genome model. A BBH pair is defined when two genes are best BLAST hits of each other with a percent identity cutoff of 60%. Majority of identified gene pairs had between 95%-100% identity (Figure S3, Table S7). This analysis was performed after each iteration of the Pan-putida model refinement. To initiate a strain-specific model, a copy of the Pan-putida model was used as the scaffold. The BBH table was used to identify genes for which there were no predicted homologs to the Pan-putida genome and were removed. This step led to the initial draft of 163 strain-specific models. FBA was then performed to simulate growth on glucose at a rate of 10 mmol/h/gCDW. For GEMs that failed the growth simulation test, gap-filling was performed to identify and add the missing essential gene(s) in individual models. Following this step, all 163 strain-specific GEMs were capable of growth on glucose. For the 24 strains with growth data, their GEMs underwent further rounds of curation. The growth ability of each strain was simulated on carbon sources provided in the Biolog and agar plate growth tests. To reconcile the difference between the simulation and the wet-lab results, automatic gap-filing^87^ was first applied followed by manual gap-filling, which focused on substrates that had large numbers of false predictions. Literature and database, e.g., Biocyc, KEGG, and Brenda, searching was performed to identified gene contents that were either missing or mis-annotated. Obtained information was then applied to the removal of false negatives or false positives. Certain false negatives were corrected by the additions of transport and exchange reactions for the carbon sources when the presence of required metabolic genes were verified. Notably, the GEM for strain 1218169.3, also known as S11^88^, had much lesser reactions (2300) compared to the average (3030) (Figure S3), possibly due to its deposited genome sequence containing 196 contigs and several frameshifted proteins. Obtained models were validated using the MEMOTE suite (https://memote.readthedocs.io/en/latest/index.html)^89^. A summary of MEMOTE reports is in Table S5. Detailed reports are provided as Supplemental Materials as well as accessible on https://doi.org/10.5281/zenodo.17382094.

### GEM refinement and curation

Using 191 carbon substrates on Biolog plates and an additional 15 aromatic carbon sources on agar plates, the 24 draft strain specific GEMs were tested for accuracy and further refinement. Since positive metabolic activities observed in Biolog experiments could result from redox reactions through the enzymatic oxidation of the substrate, cell culturing was conducted to confirm the growth of *P. putida* strains on a selected group of substrates that are of biotechnological interest as sole carbon sources, including D-glucose, D-xylose, D-galactose, L-arabinose, and formate (Figure S4A & S4B). After correlating the data, a good agreement of 74% between the Biolog and the drill-down growth assays was observed. Experimental observations of the growth assays were taken as true growth data for model constructions. Following automatic and manual gap-filling efforts^85^, the overall prediction accuracy of the 23 models improved from 64% to 91% (Figure S4C). Incorrect predictions included 7% false positives (Figure S3D). These false positives are often due to regulatory events that prevents the use of certain genes under specific conditions. Adaptive laboratory evolution can be used to overcome such regulatory interactions in some cases^90,70^. The remaining failed predictions were false negatives (Figure S4C). This failure mode results from missing metabolic capabilities that were not reconciled in the gap-filling step^87^. A large portion of the false negatives observed are accounted for by a few carbon sources, such as nucleoside bases adenosine and inosine, that supported the metabolic activities of most strains, as indicated by the horizonal white color bands in Figure S4D. The genome-scale model of the KT2440 strain, *i*JN1463, was also updated using the growth data. These strain specific model curations were incorporated into the pan-putida metabolic model which achieved an accuracy of 88% on Biolog phenotypic data and 100% on the 15 agar plate assayed aromatic carbon sources (Figure S3D)

### *In Silico* Growth Simulation

All model-based experiments were performed using Python and the COBRApy package^91^. The core biomass function from *i*JN1463 was used as the objective function for all flux balance analysis experiments. The *in silico* media was reported^32^. For growth predictions on different carbon sources the lower bound for glucose exchange was set to zero and the lower bound for the carbon source of interest was set to 10 mmol/h/gCDW.

### Pan-genome generation and Core Genome Analysis

The Cluster Database at High Identity with Tolerance, CD-HIT^39^ program with default parameters was utilized on protein sequences and annotations. Strains were first grouped according to the number of genes that they shared with the Pan-putida genome (Figure S3). Core genome began with the genome of the strain with the greatest number of genes present in the Pan-putida genome. Strains were then sequentially added, and the core genome size was recalculated based on the number of shared genes by all strains that were included.

### Allelic and Alleleome Analysis

The allelic frequency and genomic content were analyzed based on a CD-HIT scripts developed previously^92^. Briefly, CD-HIT was employed to identify shared alleles between strains based on amino acid sequences. The scripts used the Pan-putida model to associate each gene with a specific metabolic system based on the reaction(s) it catalyzed. Each allele was then linked to its corresponding gene from the Pan-putida model and the associated metabolic subsystem for that gene. Metabolic subsystems from the model were grouped into larger overarching subsystems. Alleleome analysis of the 164 strains was conducted by exporting all the alleles identified by CD-HIT and ranking the alleles with strain counts. For analysis of aromatics pathways, allele sequences of pathway enzymes were exported and manually curated to correct sequencing and annotation errors. Phylogeny was analyzed using the maximum likelihood method by IQ-TREE 2 with 500 bootstrap iterations^93, 94^. The results were visualized and edited with FigTree software version 1.4.4. (http://tree.bio.ed.ac.uk/software/figtree/).

## ACKNOWLEDGEMENT

This work was supported by Nebraska Center for Energy Science Research (to W.N.), U.S. National Institute of Health (grant P20 GM113126 to W.N. through the Nebraska Center for Integrated Biomolecular Communication), the Biorefining and Biomanufacturing program from the U.S. Department of Agriculture’s National Institute of Food and Agriculture (project award 2024-67021-42453 to W.N.), U.S. National Science Foundation Graduate Research Fellowship Program (grant 1610400 to J.M.), U.S. Department of Energy (Office of Science, Office of Biological and Environmental Research to the Joint BioEnergy Institute under contract No. DE-AC02-05CH11231).

## AUTHOR CONTRIBUTIONS

B.O.P., A.M.F., J.M.M., and W.N. designed the research. J.M., Y.H., H.V., A.A., J.K and Q.W. performed research, collected, and analyzed the data. J.M., J.K. and W.N. drafted the manuscript. J.M., W.N., A.M.F., J.M.M., J.K and B.O.P. critically revised the manuscript. All authors approved the manuscript.

## ADDITIONAL INFORMATION

**Supplementary Information** accompanies this paper is available online. Table S6 and S7 have been deposited to zenodo (https://doi.org/10.5281/zenodo.17611968) due to size limitations.

**Competing interests:** The authors declare no competing interests.

**Data Availability:** The GEMs generated in this study are made available as a supplementary file as well as at https://doi.org/10.5281/zenodo.17382094.

## REFERENCES

1. Euzeby, J. P., List of bacterial names with standing in nomenclature: a folder available on the internet. Int. J. Syst. Bacteriol. 1997, 47 (2), 590–592.

2. Palleroni, N. J., The Pseudomonas story. Environ. Microbiol. 2010, 12 (6), 1377–1383.

3. Silby, M. W.; Winstanley, C.; Godfrey, S. A. C.; Levy, S. B.; Jackson, R. W., Pseudomonas genomes: diverse and adaptable. FEMS Microbiol. Rev. 2011, 35 (4), 652–680.

4. Nikel, P. I.; Martinez-Garcia, E.; de Lorenzo, V., Biotechnological domestication of *Pseudomonads* using synthetic biology. Nat. Rev. Microbiol. 2014, 12 (5), 368–379.

5. Loeschcke, A.; Thies, S., *Pseudomonas putida* – a versatile host for the production of natural products. Appl. Microbiol. Biotechnol. 2015, 99 (15), 6197–6214.

6. Nikel, P. I.; Chavarria, M.; Danchin, A.; de Lorenzo, V., From dirt to industrial applications: *Pseudomonas putida* as a Synthetic Biology chassis for hosting harsh biochemical reactions. Curr. Opin. Chem. Biol. 2016, 34, 20–29.

7. Poblete-Castro, I.; Borrero-de Acuna, J. M.; Nikel, P. I.; Kohlstedt, M.; Wittmann, C., Host Organism: *Pseudomonas putida*. Adv. Biotechnol. 2017, 3A (Industrial Biotechnology), 299–326.

8. Nikel, P. I.; de Lorenzo, V., *Pseudomonas putida* as a functional chassis for industrial biocatalysis: From native biochemistry to trans-metabolism. Metab. Eng. 2018, 50, 142–155.

9. Martinez-Garcia, E.; de Lorenzo, V., *Pseudomonas putida* in the quest of programmable chemistry. Curr. Opin. Biotechnol. 2019, 59, 111–121.

10. Weimer, A.; Kohlstedt, M.; Volke, D. C.; Nikel, P. I.; Wittmann, C., Industrial biotechnology of *Pseudomonas putida*: advances and prospects. Appl. Microbiol. Biotechnol. 2020, 104 (18), 7745–7766.

11. Nogales, J.; García, J. L.; Díaz, E., Degradation of Aromatic Compounds in Pseudomonas: A Systems Biology View. In Aerobic Utilization of Hydrocarbons, Oils and Lipids, Rojo, F., Ed. Springer International Publishing: Cham, 2017; pp 1–49.

12. Dvorak, P.; Nikel, P. I.; Damborsky, J.; de Lorenzo, V., Bioremediation 3.0: Engineering pollutant-removing bacteria in the times of systemic biology. Biotechnol. Adv. 2017, 35 (7), 845–866.

13. Martinez-Garcia, E.; de Lorenzo, V., Molecular tools and emerging strategies for deep genetic/genomic refactoring of Pseudomonas. Curr. Opin. Biotechnol. 2017, 47, 120–132.

14. Martin-Pascual, M.; Batianis, C.; Bruinsma, L.; Asin-Garcia, E.; Garcia-Morales, L.; Weusthuis, R. A.; van Kranenburg, R.; Martins dos Santos, V. A. P., A navigation guide of synthetic biology tools for *Pseudomonas putida*. Biotechnol. Adv. 2021, 49, 107732.

15. He, X.; Gao, T.; Chen, Y.; Liu, K.; Guo, J.; Niu, W., Genetic Code Expansion in *Pseudomonas putida* KT2440. ACS Synth. Biol. 2022, 11 (11), 3724–3732.

16. Kozaeva, E.; Volkova, S.; Matos, M. R. A.; Mezzina, M. P.; Wulff, T.; Volke, D. C.; Nielsen, L. K.; Nikel, P. I., Model-guided dynamic control of essential metabolic nodes boosts acetyl-coenzyme A-dependent bioproduction in rewired *Pseudomonas putida*. Metab. Eng. 2021, 67, 373–386.

17. Lim, H. G.; Rychel, K.; Sastry, A. V.; Bentley, G. J.; Mueller, J.; Schindel, H. S.; Larsen, P. E.; Laible, P. D.; Guss, A. M.; Niu, W.; Johnson, C. W.; Beckham, G. T.; Feist, A. M.; Palsson, B. O., Machine-learning from *Pseudomonas putida* KT2440 transcriptomes reveals its transcriptional regulatory network. Metab. Eng. 2022, 72, 297–310.

18. Edwards, J. S.; Palsson, B. O., Systems properties of the *Haemophilus influenzae*Rd metabolic genotype. J. Biol. Chem. 1999, 274 (25), 17410–17416.

19. Price, N. D.; Papin, J. A.; Schilling, C. H.; Palsson, B. O., Genome-scale microbial in silico models: the constraints-based approach. Trends Biotechnol. 2003, 21 (4), 162–169.

20. Thiele, I.; Palsson, B. O., A protocol for generating a high-quality genome-scale metabolic reconstruction. Nat. Protoc. 2010, 5 (1), 93–121.

21. O’Brien, E. J.; Monk, J. M.; Palsson, B. O., Using genome-scale models to predict biological capabilities. Cell 2015, 161 (5), 971–987.

22. Gu, C.; Kim, G. B.; Kim, W. J.; Lee, S. Y.; Kim, H. U.; Kim, H. U.; Lee, S. Y.; Kim, H. U.; Lee, S. Y., Current status and applications of genome-scale metabolic models. Genome Biol. 2019, 20 (1), 121.

23. Monk, J. M.; Charusanti, P.; Aziz, R. K.; Lerman, J. A.; Premyodhin, N.; Orth, J. D.; Feist, A. M.; Palsson, B. O., Genome-scale metabolic reconstructions of multiple *Escherichia coli* strains highlight strain-specific adaptations to nutritional environments. Proc. Natl. Acad. Sci. U. S. A. 2013, 110 (50), 20338–20343.

24. Bosi, E.; Monk, J. M.; Aziz, R. K.; Fondi, M.; Nizet, V.; Palsson, B. O., Comparative genome-scale modelling of *Staphylococcus aureus* strains identifies strain-specific metabolic capabilities linked to pathogenicity. Proc. Natl. Acad. Sci. U. S. A. 2016, 113 (26), E3801–E3809.

25. Seif, Y.; Kavvas, E.; Lachance, J.-C.; Yurkovich, J. T.; Nuccio, S.-P.; Fang, X.; Catoiu, E.; Raffatellu, M.; Palsson, B. O.; Monk, J. M., Genome-scale metabolic reconstructions of multiple *Salmonella* strains reveal serovar-specific metabolic traits. Nat. Commun. 2018, 9 (1), 3771.

26. Norsigian, C. J.; Danhof, H. A.; Brand, C. K.; Midani, F. S.; Broddrick, J. T.; Savidge, T. C.; Britton, R. A.; Palsson, B. O.; Spinler, J. K.; Monk, J. M., Systems biology approach to functionally assess the *Clostridioides difficile* pangenome reveals genetic diversity with discriminatory power. Proc. Natl. Acad. Sci. U. S. A. 2022, 119 (18), e2119396119.

27. Nelson, K. E.; Weinel, C.; Paulsen, I. T.; Dodson, R. J.; Hilbert, H.; Martins dos Santos, V. A. P.; Fouts, D. E.; Gill, S. R.; Pop, M.; Holmes, M.; Brinkac, L.; Beanan, M.; DeBoy, R. T.; Daugherty, S.; Kolonay, J.; Madupu, R.; Nelson, W.; White, O.; Peterson, J.; Khouri, H.; Hance, I.; Lee, P. C.; Holtzapple, E.; Scanlan, D.; Tran, K.; Moazzez, A.; Utterback, T.; Rizzo, M.; Lee, K.; Kosack, D.; Moestl, D.; Wedler, H.; Lauber, J.; Stjepandic, D.; Hoheisel, J.; Straetz, M.; Heim, S.; Kiewitz, C.; Eisen, J.; Timmis, K. N.; Dusterhoft, A.; Tummler, B.; Fraser, C. M., Complete genome sequence and comparative analysis of the metabolically versatile Pseudomonas putida KT2440. Environ. Microbiol. 2002, 4 (12), 799–808.

28. Belda, E.; van Heck, R. G. A.; Jose Lopez-Sanchez, M.; Cruveiller, S.; Barbe, V.; Fraser, C.; Klenk, H.-P.; Petersen, J.; Morgat, A.; Nikel, P. I.; Vallenet, D.; Rouy, Z.; Sekowska, A.; Martins dos Santos, V. A. P.; de Lorenzo, V.; Danchin, A.; Medigue, C., The revisited genome of Pseudomonas putida KT2440 enlightens its value as a robust metabolic chassis. Environ. Microbiol. 2016, 18 (10), 3403–3424.

29. Kampers, L. F.; Volkers, R. J.; Martins Dos Santos, V. A., Pseudomonas putida KT 2440 is HV 1 certified, not GRAS. Microb. Biotechnol. 2019, 12 (5), 845–848.

30. Nogales, J.; Palsson, B. O.; Thiele, I., A genome-scale metabolic reconstruction of *Pseudomonas putida* KT2440: iJN746 as a cell factory. BMC Syst. Biol. 2008, 2, 79.

31. Puchalka, J.; Oberhardt, M. A.; Godinho, M.; Bielecka, A.; Regenhardt, D.; Timmis, K. N.; Papin, J. A.; Martins dos Santos, V. A. P., Genome-scale reconstruction and analysis of the Pseudomonas putida KT2440 metabolic network facilitates applications in biotechnology. PLoS Comput. Biol. 2008, 4 (10), e1000210.

32. Nogales, J.; Mueller, J.; Gudmundsson, S.; Canalejo, F. J.; Duque, E.; Monk, J.; Feist, A. M.; Ramos, J. L.; Niu, W.; Palsson, B. O., High-quality genome-scale metabolic modelling of *Pseudomonas putida* highlights its broad metabolic capabilities. Environ. Microbiol. 2020, 22 (1), 255–269.

33. Yonezuka, K.; Shimodaira, J.; Tabata, M.; Ohji, S.; Hosoyama, A.; Kasai, D.; Yamazoe, A.; Fujita, N.; Ezaki, T.; Fukuda, M., Phylogenetic analysis reveals the taxonomically diverse distribution of the *Pseudomonas putida* group. J. Gen. Appl. Microbiol. 2017, 63 (1), 1–10.

34. Shea, A.; Wolcott, M.; Daefler, S.; Rozak, D. A., Biolog phenotype microarrays. Methods Mol. Biol. 2012, 881, 331–373.

35. Schellenberger, J.; Park, J. O.; Conrad, T. M.; Palsson, B. O., BiGG: a Biochemical Genetic and Genomic knowledgebase of large scale metabolic reconstructions. BMC Bioinf. 2010, 11.

36. Becker, J.; Wittmann, C., A field of dreams: Lignin valorization into chemicals, materials, fuels, and health-care products. Biotechnol. Adv. 2019, 37 (6), 107360.

37. Bugg, T. D. H.; Williamson, J. J.; Alberti, F., Microbial hosts for metabolic engineering of lignin bioconversion to renewable chemicals. Renewable Sustainable Energy Rev. 2021, 152, 111674.

38. Krishnan, K. J., Hefner, Y., Szubin, R., Monk, J., Pride, D. T., & Palsson, B. (2025). PMkbase (version 1.0): an interactive web-based tool for tracking bacterial metabolic traits using phenotype microarrays made interoperable with sequence information and visualizing/processing PM data. *Microbiology Spectrum*, e03279–24.

39. Huang, Y.; Niu, B.; Gao, Y.; Fu, L.; Li, W., CD-HIT Suite: a web server for clustering and comparing biological sequences. Bioinformatics 2010, 26 (5), 680–682.

40. Inglin, R. C.; Meile, L.; Stevens, M. J. A., Clustering of pan– and core-genome of *Lactobacillus* provides novel evolutionary insights for differentiation. BMC Genomics 2018, 19, 284.

41. Passarelli-Araujo, H.; Jacobs, S. H.; Franco, G. R.; Venancio, T. M., Phylogenetic analysis and population structure of *Pseudomonas alloputida*. Genomics 2021, 113 (6), 3762–3773.

42. Catoiu, E. A.; Phaneuf, P.; Monk, J.; Palsson, B. O., Whole-genome sequences from wild-type and laboratory-evolved strains define the alleleome and establish its hallmarks. Proc. Natl. Acad. Sci. U. S. A. 2023, 120 (15), e2218835120.

43. Moreno, R.; Ruiz-Manzano, A.; Yuste, L.; Rojo, F., The Pseudomonas putida Crc global regulator is an RNA binding protein that inhibits translation of the AlkS transcriptional regulator. Mol. Microbiol. 2007, 64 (3), 665–675.

44. Romeo, T., Global regulation by the small RNA-binding protein CsrA and the non-coding RNA molecule CsrB. Mol. Microbiol. 1998, 29 (6), 1321–1330.

45. Rojo, F., Carbon catabolite repression in Pseudomonas: optimizing metabolic versatility and interactions with the environment. FEMS Microbiol. Rev. 2010, 34 (5), 658–684.

46. Yuste, L.; Hervas, A. B.; Canosa, I.; Tobes, R.; Jimenez, J. I.; Nogales, J.; Perez-Perez, M. M.; Santero, E.; Diaz, E.; Ramos, J.-L.; de Lorenzo, V.; Rojo, F., Growth phase-dependent expression of the Pseudomonas putida KT2440 transcriptional machinery analysed with a genome-wide DNA microarray. Environ. Microbiol. 2006, 8 (1), 165–177.

47. Ijaq, J.; Chandra, D.; Ray, M. K.; Jagannadham, M. V., Investigating the functional role of hypothetical proteins from an antarctic bacterium Pseudomonas sp. Lz4W: emphasis on identifying proteins involved in cold adaptation. Front. Genet. 2022, 13, 825269.

48. Silhavy, T. J.; Kahne, D.; Walker, S., The bacterial cell envelope. Cold Spring Harbor Perspect. Biol. 2010, 2 (5).

49. Feist, A. M.; Herrgard, M. J.; Thiele, I.; Reed, J. L.; Palsson, B. O., Reconstruction of biochemical networks in microorganisms. Nat. Rev. Microbiol. 2009, 7 (2), 129–143.

50. Olson, R. D.; Assaf, R.; Brettin, T.; Conrad, N.; Cucinell, C.; Davis, J. J.; Dempsey, D. M.; Dickerman, A.; Dietrich, E. M.; Kenyon, R. W.; Kuscuoglu, M.; Lefkowitz, E. J.; Lu, J.; Machi, D.; Macken, C.; Mao, C.; Niewiadomska, A.; Nguyen, M.; Olsen, G. J.; Overbeek, J. C.; Parrello, B.; Parrello, V.; Porter, J. S.; Pusch, G. D.; Shukla, M.; Singh, I.; Stewart, L.; Tan, G.; Thomas, C.; VanOeffelen, M.; Vonstein, V.; Wallace, Z. S.; Warren, A. S.; Wattam, A. R.; Xia, F.; Yoo, H.; Zhang, Y.; Zmasek, C. M.; Scheuermann, R. H.; Stevens, R. L., Introducing the bacterial and viral bioinformatics resource center (BV-BRC): a resource combining PATRIC, IRD and ViPR. Nucleic Acids Res. 2023, 51 (D1), D678.

51. Guzman, G. I.; Olson, C. A.; Hefner, Y.; Phaneuf, P. V.; Catoiu, E.; Crepaldi, L. B.; Micas, L. G.; Palsson, B. O.; Feist, A. M., Reframing gene essentiality in terms of adaptive flexibility. BMC Syst. Biol. 2018, 12, 143.

52. Sandberg, T. E.; Salazar, M. J.; Weng, L. L.; Palsson, B. O.; Feist, A. M., The emergence of adaptive laboratory evolution as an efficient tool for biological discovery and industrial biotechnology. Metab. Eng. 2019, 56, 1–16.

53. Bruinsma, L.; Wenk, S.; Claassens, N. J.; Martins dos Santos, V. A. P., Paving the way for synthetic C1 – Metabolism in Pseudomonas putida through the reductive glycine pathway. Metab. Eng. 2023, 76, 215–224.

54. Bochner, B. R., Global phenotypic characterization of bacteria. FEMS Microbiol. Rev. 2009, 33 (1), 191–205.

55. Roca, A.; Rodriguez-Herva, J. J.; Ramos, J. L., Redundancy of enzymes for formaldehyde detoxification in Pseudomonas putida. J. Bacteriol. 2009, 191 (10), 3367–3374.

56. Fuchs, G.; Boll, M.; Heider, J., Microbial degradation of aromatic compounds – from one strategy to four. Nat. Rev. Microbiol. 2011, 9 (11), 803–816.

57. Wells, T.; Ragauskas, A. J., Biotechnological opportunities with the β-ketoadipate pathway. Trends Biotechnol. 2012, 30 (12), 627–637.

58. Luengo, J. M.; Garcia, J. L.; Olivera, E. R., The phenylacetyl-CoA catabolon: a complex catabolic unit with broad biotechnological applications. Mol. Microbiol. 2001, 39 (6), 1434–1442.

59. Arias-Barrau, E.; Olivera, E. R.; Luengo, J. M.; Fernandez, C.; Galan, B.; Garcia, J. L.; Diaz, E.; Minambres, B., The homogentisate pathway: A central catabolic pathway involved in the degradation of L-phenylalanine, L-tyrosine, and 3-hydroxyphenylacetate in Pseudomonas putida. J. Bacteriol. 2004, 186 (15), 5062–5077.

60. Arcos, M.; Olivera, E. R.; Arias, S.; Naharro, G.; Luengo, J. M., The 3,4-dihydroxyphenylacetic acid catabolon, a catabolic unit for degradation of biogenic amines tyramine and dopamine in Pseudomonas putida U. Environ. Microbiol. 2010, 12 (6), 1684–1704.

61. 61.

62. Luengo, J. M.; Olivera, E. R., Catabolism of biogenic amines in Pseudomonas species. Environ. Microbiol. 2020, 22 (4), 1174–1192.

63. Shingler, V.; Powlowski, J.; Marklund, U., Nucleotide sequence and functional analysis of the complete phenol/3,4-dimethylphenol catabolic pathway of Pseudomonas sp. strain CF600. J. Bacteriol. 1992, 174 (3), 711.

64. Powlowski, J.; Shingler, V., Genetics and biochemistry of phenol degradation by Pseudomonas sp. CF600. Biodegradation 1994, 5 (3-4), 219.

65. Chen, Y.-F.; Chao, H.; Zhou, N.-Y., The catabolism of 2,4-xylenol and p-cresol share the enzymes for the oxidation of para-methyl group in Pseudomonas putida NCIMB 9866. Appl. Microbiol. Biotechnol. 2014, 98 (3), 1349–1356.

66. Dvorak, P.; de Lorenzo, V., Refactoring the upper sugar metabolism of Pseudomonas putida for co-utilization of cellobiose, xylose, and glucose. Metab. Eng. 2018, 48, 94–108.

67. Peabody, V. G. L.; Elmore, J. R.; Martinez-Baird, J.; Guss, A. M., Engineered Pseudomonas putida KT2440 co-utilizes galactose and glucose. Biotechnol. Biofuels 2019, 12 (1), 295.

68. Elmore, J. R.; Dexter, G. N.; Salvachua, D.; O’Brien, M.; Klingeman, D. M.; Gorday, K.; Michener, J. K.; Peterson, D. J.; Beckham, G. T.; Guss, A. M., Engineered Pseudomonas putida simultaneously catabolizes five major components of corn stover lignocellulose: Glucose, xylose, arabinose, p-coumaric acid, and acetic acid. Metab. Eng. 2020, 62, 62–71.

69. Ling, C.; Peabody, G. L.; Salvachua, D.; Kim, Y.-M.; Kneucker, C. M.; Calvey, C. H.; Monninger, M. A.; Munoz, N. M.; Poirier, B. C.; Ramirez, K. J.; St. John, P. C.; Woodworth, S. P.; Magnuson, J. K.; Burnum-Johnson, K. E.; Guss, A. M.; Johnson, C. W.; Beckham, G. T., Muconic acid production from glucose and xylose in Pseudomonas putida via evolution and metabolic engineering. Nat. Commun. 2022, 13 (1), 4925.

70. Bujdos, D.; Popelarova, B.; Volke, D. C.; Nikel, P. I.; Sonnenschein, N.; Dvorak, P., Engineering of Pseudomonas putida for accelerated co-utilization of glucose and cellobiose yields aerobic overproduction of pyruvate explained by an upgraded metabolic model. Metab. Eng. 2023, 75, 29–46.

71. Lim, H. G.; Eng, T.; Banerjee, D.; Alarcon, G.; Lau, A. K.; Park, M.-R.; Simmons, B. A.; Palsson, B. O.; Singer, S. W.; Mukhopadhyay, A.; Feist, A. M., Generation of Pseudomonas putida kt2440 strains with efficient utilization of xylose and galactose via adaptive lab. evolution. ACS Sustainable Chem. Eng. 2021, 9 (34), 11512–11523.

72. Jimenez, J. I.; Minambres, B.; Garcia, J. L.; Diaz, E., Genomic analysis of the aromatic catabolic pathways from Pseudomonas putida KT2440. Environ. Microbiol. 2002, 4 (12), 824–841.

73. Notonier, S.; Werner, A. Z.; Kuatsjah, E.; Dumalo, L.; Abraham, P. E.; Hatmaker, E. A.; Hoyt, C. B.; Amore, A.; Ramirez, K. J.; Woodworth, S. P.; Klingeman, D. M.; Giannone, R. J.; Guss, A. M.; Hettich, R. L.; Eltis, L. D.; Johnson, C. W.; Beckham, G. T., Metabolism of syringyl lignin-derived compounds in Pseudomonas putida enables convergent production of 2-pyrone-4,6-dicarboxylic acid. Metab. Eng. 2021, 65, 111–122.

74. Mueller, J.; Willett, H.; Feist, A. M.; Niu, W., Engineering Pseudomonas putida for improved utilization of syringyl aromatics. Biotechnol. Bioeng. 2022, 119 (9), 2541–2550.

75. Niu, W.; Kramer, L.; Mueller, J.; Liu, K.; Guo, J., Metabolic engineering of Escherichia coli for the de novo stereospecific biosynthesis of 1,2-propanediol through lactic acid. Metab. Eng. Commun. 2019, 8, e00082.

76. Niu, W.; Willett, H.; Mueller, J.; He, X.; Kramer, L.; Ma, B.; Guo, J., Direct biosynthesis of adipic acid from lignin-derived aromatics using engineered Pseudomonas putida KT2440. Metab. Eng. 2020, 59, 151–161.

77. Wick, R. R.; Judd, L. M.; Gorrie, C. L.; Holt, K. E., Unicycler: Resolving bacterial genome assemblies from short and long sequencing reads. PLoS Comput. Biol. 2017, 13 (6), e1005595/1-e1005595/22.

78. Seemann, T., Prokka: rapid prokaryotic genome annotation. Bioinformatics 2014, 30 (14), 2068–2069.

79. Gurevich, A.; Saveliev, V.; Vyahhi, N.; Tesler, G., QUAST: quality assessment tool for genome assemblies. Bioinformatics 2013, 29 (8), 1072–1075.

80. Tegenfeldt, F.; Kuznetsov, D.; Manni, M.; Berkeley, M.; Zdobnov, E. M.; Kriventseva, E. V., OrthoDB and BUSCO update: annotation of orthologs with wider sampling of genomes. Nucleic Acids Res 2025, 53 (D1), D516–D522.

81. Edgar, R. C., MUSCLE: a multiple sequence alignment method with reduced time and space complexity. BMC Bioinf. 2004, 5, 113.

82. Kumar, S.; Stecher, G.; Li, M.; Knyaz, C.; Tamura, K., MEGA X: molecular evolutionary genetics analysis across computing platforms. Mol. Biol. Evol. 2018, 35 (6), 1547–1549.

83. Tamura, K.; Nei, M., Estimation of the number of nucleotide substitutions in the control region of mitochondrial DNA in humans and chimpanzees. Mol. Biol. Evol. 1993, 10 (3), 512–526.

84. Monk, J.; Nogales, J.; Palsson, B. O., Optimizing genome-scale network reconstructions. Nat. Biotechnol. 2014, 32 (5), 447–452.

85. Davis, J. J.; Wattam, A. R.; Aziz, R. K.; Brettin, T.; Butler, R.; Butler, R. M.; Chlenski, P.; Conrad, N.; Dickerman, A.; Dietrich, E. M.; Gabbard, J. L.; Gerdes, S.; Guard, A.; Kenyon, R. W.; Machi, D.; Mao, C.; Murphy-Olson, D.; Nguyen, M.; Nordberg, E. K.; Olsen, G. J.; Olson, R. D.; Overbeek, J. C.; Overbeek, R.; Parrello, B.; Pusch, G. D.; Shukla, M.; Thomas, C.; Vanoeffelen, M.; Vonstein, V.; Warren, A. S.; Xia, F.; Xie, D.; Yoo, H.; Stevens, R., The PATRIC bioinformatics resource center: expanding data and analysis capabilities. Nucleic Acids Res. 2020, 48 (D1), D606–D612.

86. Norsigian, C. J.; Fang, X.; Seif, Y.; Monk, J. M.; Palsson, B. O., A workflow for generating multi-strain genome-scale metabolic models of prokaryotes. Nat. Protoc. 2020, 15 (1), 1–14.

87. Schmelling, N., Reciprocal Best Hit BLAST v2. protocols.io 2018, 10.17504/protocols.io.q3rdym6.

88. Reed, J. L.; Patel, T. R.; Chen, K. H.; Joyce, A. R.; Applebee, M. K.; Herring, C. D.; Bui, O. T.; Knight, E. M.; Fong, S. S.; Palsson, B. O., Systems approach to refining genome annotation. Proc. Natl. Acad. Sci. U. S. A. 2006, 103 (46), 17480–17484.

89. Ponraj, P.; Shankar, M.; Ilakkiam, D.; Rajendhran, J.; Gunasekaran, P., Genome sequence of the plant growth-promoting rhizobacterium *Pseudomonas putida* S11. J. Bacteriol. 2012, 194 (21), 6015.

90. Lieven, C.; Beber, M. E.; Olivier, B. G.; Bergmann, F. T.; Ataman, M.; Babaei, P.; Bartell, J. A.; Blank, L. M.; Chauhan, S.; Correia, K.; Diener, C.; Drager, A.; Ebert, B. E.; Edirisinghe, J. N.; Faria, J. P.; Feist, A. M.; Fengos, G.; Fleming, R. M. T.; Garcia-Jimenez, B.; Hatzimanikatis, V.; van Helvoirt, W.; Henry, C. S.; Hermjakob, H.; Herrgaard, M. J.; Kaafarani, A.; Kim, H. U.; King, Z.; Klamt, S.; Klipp, E.; Koehorst, J. J.; Konig, M.; Lakshmanan, M.; Lee, D.-Y.; Lee, S. Y.; Lee, S.; Lewis, N. E.; Liu, F.; Ma, H.; Machado, D.; Mahadevan, R.; Maia, P.; Mardinoglu, A.; Medlock, G. L.; Monk, J. M.; Nielsen, J.; Nielsen, L. K.; Nogales, J.; Nookaew, I.; Palsson, B. O.; Papin, J. A.; Patil, K. R.; Poolman, M.; Price, N. D.; Resendis-Antonio, O.; Richelle, A.; Rocha, I.; Sanchez, B. J.; Schaap, P. J.; Malik Sheriff, R. S.; Shoaie, S.; Sonnenschein, N.; Teusink, B.; Vilaca, P.; Vik, J. O.; Wodke, J. A. H.; Xavier, J. C.; Yuan, Q.; Zakhartsev, M.; Zhang, C., MEMOTE for standardized genome-scale metabolic model testing. Nat. Biotechnol. 2020, 38 (3), 272–276.

91. Guzman, G. I.; Sandberg, T. E.; LaCroix, R. A.; Nyerges, A.; Papp, H.; de Raad, M.; King, Z. A.; Hefner, Y.; Northen, T. R.; Notebaart, R. A.; Pal, C.; Palsson, B. O.; Papp, B.; Feist, A. M., Enzyme promiscuity shapes adaptation to novel growth substrates. Mol. Syst. Biol. 2019, 15 (4).

92. Ebrahim, A.; Lerman, J. A.; Palsson, B. O.; Hyduke, D. R., COBRApy: COnstraints-Based Reconstruction and Analysis for Python. BMC Systems Biology 2013, 7 (1), 74.

93. Hyun, J. C.; Monk, J. M.; Palsson, B. O., Comparative pangenomics: analysis of 12 microbial pathogen pangenomes reveals conserved global structures of genetic and functional diversity. BMC Genomics 2022, 23 (1), 7.

94. Nguyen, L.-T.; Schmidt, H. A.; von Haeseler, A.; Minh, B. Q., IQ-TREE: a fast and effective stochastic algorithm for estimating maximum-likelihood phylogenies. Mol. Biol. Evol. 2015, 32 (1), 268–274.

95. Minh, B. Q.; Schmidt, H. A.; Chernomor, O.; Schrempf, D.; Woodhams, M. D.; Von Haeseler, A.; Lanfear, R., IQ-TREE 2: new models and efficient methods for phylogenetic inference in the genomic era. Mol. Biol. Evol. 2020, 37 (5), 1530–1534.

